# Structural basis for the enhanced infectivity and immune evasion of Omicron subvariants

**DOI:** 10.1101/2022.07.13.499586

**Authors:** Yaning Li, Yaping Shen, Yuanyuan Zhang, Renhong Yan

**Affiliations:** Center for Infectious Disease Research, Westlake Laboratory of Life Sciences and Biomedicine, Key Laboratory of Structural Biology of Zhejiang Province, School of Life Sciences, Westlake University, Hangzhou 310024, Zhejiang Province, China; Beijing Advanced Innovation Center for Structural Biology, Tsinghua-Peking Joint Center for Life Sciences, School of Life Sciences, Tsinghua University, Beijing 100084, China

## Abstract

The Omicron variants of SARS-CoV-2 have recently become the globally dominant variants of concern in the COVID-19 pandemic. At least five major Omicron sub-lineages have been characterized: BA.1, BA.2, BA.3, BA.4 and BA.5. They all possess over 30 mutations on the Spike (S) protein. Here we report the cryo-EM structures of the trimeric S proteins from the five subvariants, of which BA.4 and BA.5 share the same mutations of S protein, each in complex with the surface receptor ACE2. All three receptor binding domains of S protein from BA.2 and BA.4/BA.5 are “up”, while the BA.1 S protein has two “up” and one “down”. The BA.3 S protein displays increased heterogeneity, with the majority in the all “up” RBD state. The differentially preferred conformations of the S protein are consistent with their varied transmissibilities. Analysis of the well defined S309 and S2K146 epitopes reveals the underlie immune evasion mechanism of Omicron subvariants.

## Introduction

The ongoing emergence of SARS-CoV-2 variants, especially the recent variant of concern (VOC) Omicron, continues to impose health threat due to their enhanced transmission and immune evasion(*1-3*). The Omicron has developed into at least five major sub-lineages: BA.1 (B.1.1.529.1), BA.2 (B.1.1.529.2), BA.3 (B.1.1.529.3), BA.4, and BA.5. The BA.1 sub-lineage has quickly outcompeted the Delta variant across the world since the end of 2021^ref.^(*4*). Two months later, the BA.2 subvariant, with its increased transmissibility, has replaced the BA.2 and spread worldwide(*1*). In recent days, the BA.4/BA.5 sublineages, which share the same S protein mutations (hereafter referred to as BA.4), were reported in Botswana and South Africa and displayed a higher transmission advantage than BA.2 subvariant(*5*). Additionally, the BA.3 sub-lineage is at a low prevalence for the time being but continues to spread(*6*).

The five sub-lineages of Omicron all contain multiple site changes in the Spike (S) protein, which mediates receptor recognition and facilitates membrane fusion with the host cells(*7, 8*). BA.1, BA.2, BA.3, and BA.4/BA.5 contain 37, 31, 33, and 34 mutations, respectively, in the S protein compared to the original strain (hereafter referred to as the WT strain) (Fig. 1a). These mutations potentially strengthen the variants’ transmission capability(*7*). Among these, 15 mutations in BA.1 are located on the receptor-binding domain (RBD), which is responsible for direct association with angiotensin-converting enzyme 2 (ACE2), the surface receptor on the host cells(*9, 10*). In addition to 12 common mutations with BA.1, BA.2 has 4 distinct ones, S371F, T376A, D405N, and R408S, on the RBD. BA.4 might evolved from BA.2 and has 3 additional mutations, L452R, F486V, and R493Q, on the RBD. Besides, BA.3 has combined mutations from BA.1 and BA.2 (Fig. 1a). Most recently, there emerged a new sub-lineage, Omicron XE, recombined from BA.1 and BA.2 within the nonstructural protein (NSP) 6 of the SARS-CoV-2 genome. Omicron XE shares identical S protein with BA.2, but its recorded transmission rate is 9.8% higher than BA.2^ref.^(*11*).

**Fig. 1:**
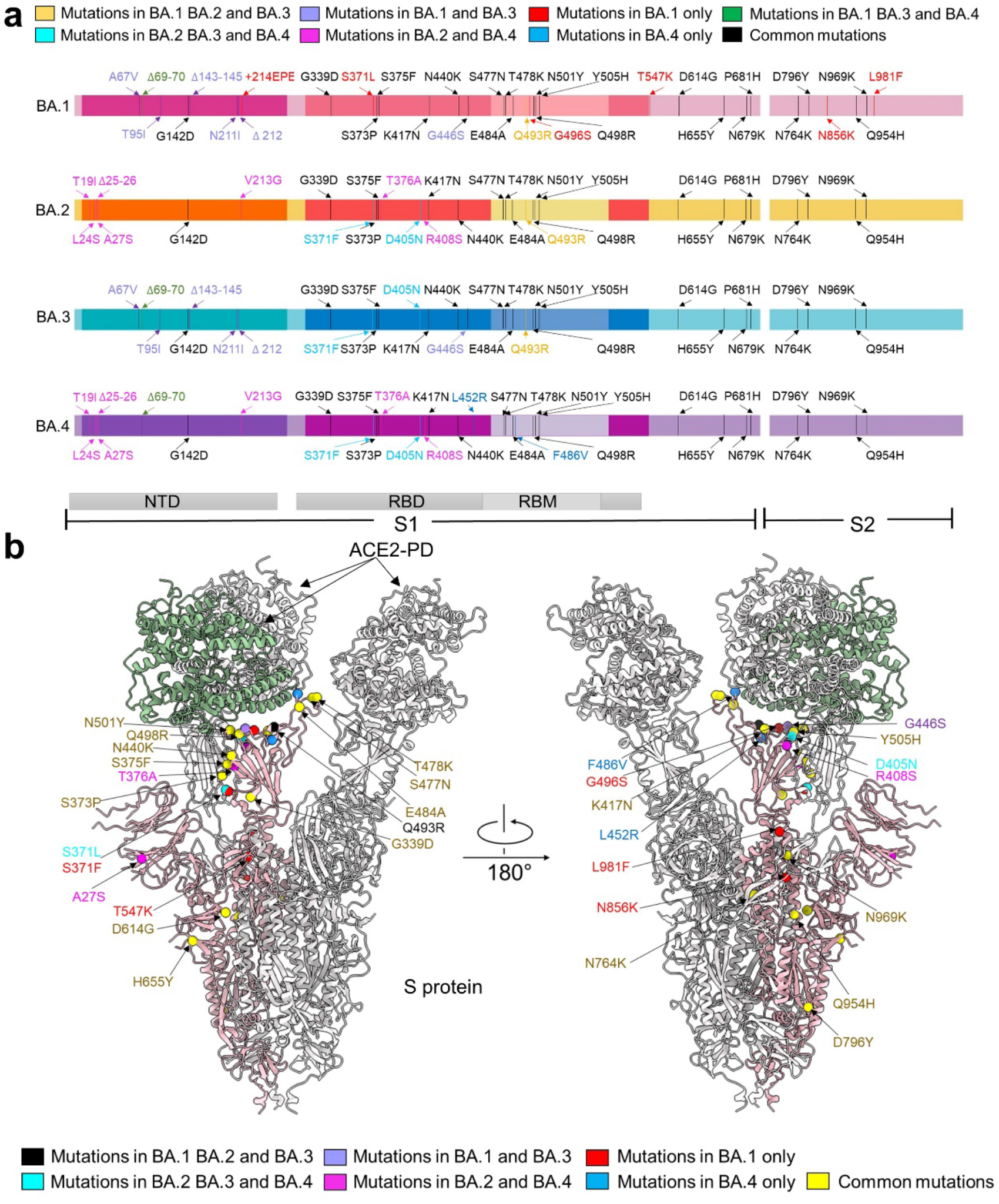
The distribution of mutations of S protein from Omicron sub-lineages. **a**, Summary of mutations mapped to the extracellular domain of the Spike protein (S-ECD) from the four Omicron sub-lineages, BA.1, BA.2, BA.3 and BA.4. They share 20 common mutations, which are labeled black. Mutations common to BA.1/2/3, BA.2/3/4 or BA.1/3/4 are colored yellow, cyan or green, respectively. The 8 mutations in BA.1 and BA.3, but not BA.2 and BA.4, are colored purple and the mutations only shared by BA.2 and BA.4 are labeled magenta.The unique mutations in BA.1 and BA.4 are colored red and blue, respectively. NTD: amino terminal domain. RBD: receptor binding domain. RBM: receptor-binding motif. **b**, Mapping of mutations to the S protein of Omicron subvariants. The common mutations of all four subvariants are colored yellow. Mutations common to BA.1, BA.2 and BA.3 are colored black. Mutations shared by BA.2, BA.3 and BA.4 are colored cyan. The mutations in BA.1 and BA.3, but not BA.2 and BA.4, are colored purple and the mutations only shared by BA.2 and BA.4 are colored magenta.The unique mutations in BA.1 and BA.4 are colored red and blue, respectively.

Despite the large number of mutations, all these Omicron subvariants still exploit ACE2 as the host receptor(*12-14*). For SARS-CoV-2 infection, the S protein is cleaved into the S1 and S2 subunits by the host furin protease(*15-19*). S1 subunit contains the N-terminal domain (NTD) and RBD. The RBD may exhibit “up” and “down” conformations in reference to the viral surface, and only the “up” RBD can expose the ACE2 binding site (*9*). Binding of the RBD to the peptidase domain (PD) of ACE2 triggers the conformational changes of the trimeric S protein, which further exposes the fusion peptide in S2 to facilitate membrane fusion with host cells(*9, 20-22*).

Omicron sub-lineages show increased immune escape capability(*23*). BA.1 was reported to evade neutralization induced by various vaccines or infection with other SARS-CoV-2 variants(*24-26*). Some patients who recovered from BA.1 were then infected by BA.2 within a short period(*27, 28*), suggesting that the antibodies generated from the early Omicron BA.1 infection might fail to neutralize BA.2. Additionally, BA.4/5 exhibits compromised neutralization to the sera from triple AstraZeneca or Pfizer vaccinated individuals than BA.1 and BA.2 subvariants(*29*). Indeed, the BA.2 and BA.4 subvariant can escape from several neutralizing antibodies, including S309 (sotrovimab, short as S309) that can effectively neutralize Alpha, Beta, Gamma, Delta, and BA.1 of SARS-CoV-2^refs.^(*12, 23*) and S2K146 that could mediate broad sarbecovirus neutralization including BA.2 and BA.3, while largely compromised the neutralization to BA.4^refs.^(*30, 31*). Fortunately, the recently authorized LY-CoV1404 (bebtelovimab) remains potent in neutralizing all Omicron sub-lineages(*23, 31, 32*).

The mutations on the S protein of different Omicron subvariants may underlie their altered properties for receptor recognition, transmission, and immune escape. It is thus critical to pinpoint the specific mutations on the S protein that are responsible for these changes. Structures of BA.1 S protein in complex with ACE2 were reported (*13, 33, 34*). However, these structures cannot explain the enhanced infectivity and acquired antibody resistance of BA.2, BA.3, and BA.4^ref.^(*6*).

To address this important question, we set out to systematically examine structures of ACE2-bound S protein from BA.1, BA.2, BA.3, and BA.4. We show that all three RBDs of the trimeric S protein from BA.2 and BA.4 are “up”, while BA.1 has two “up” and one “down”. Although the majority of the BA.3 S proteins have three “up” RBDs, a small portion has two “up” and one “down”. This analysis immediately affords an explanation for their different infectivities, as the up RBDs are to be recognized by ACE2. We also show that the shift of glycosylation moieties may lead to weakened neutralization of S309, and two mutations, L452R and F486V, of BA.4 decrease the binding to S2K146.

## Results

### Biochemical characterization and structural determination of the complex formation between Omicron Spike proteins and ACE2-PD

To investigate the biochemical characteristics of Omicron subvariants, binding affinity between their RBDs and ACE2-PD was measured and compared. The monomeric human ACE2-PD binds to the RBD of Omicron BA.1/2/3/4 with KD of 11.50±0.02 nM, 4.15±0.01 nM, 6.57±0.02 nM, and 4.55±0.02 nM, respectively, approximately 2-4 folds higher than that of WT-RBD (18.40±0.02 nM) (Supplementary Fig. 1). Our measured affinities of WT and BA.1 with ACE2-PD are similar to a previous report (*35*). Although the enhanced affinity with ACE2 by the Omicron variants may in part account for the increased transmissibility of Omicron variants, it cannot explain the particularly higher infectivity of BA.4 than BA.2 and BA.3. We then employed single-particle cryo-EM to solve the structures of the three trimeric S proteins each in complex with ACE2-PD.

We have previously solved the structure of S protein in BA.1 bound with ACE2-PD at an overall resolution of 3.3 Å(*36*). Identical protocol was used for protein expression, purification, and cryo-sample preparation for the three complexes. Please refer to Methods for details. The structures of ACE2-PD bound trimeric S protein from BA.2, BA.3, and BA.4 were determined at overall resolutions of 3.3 Å, 3.4 Å, and 2.8 Å, respectively (Supplementary Figs. 2-7 and Supplementary Table 1). For simplicity, we will refer to the three complexes as BA.1/2/3/4-SA.

The structures allow mapping of Omicron mutations, which are mainly distributed on the surface of the S protein (Fig. 1b). Mutations on the spike glycoprotein of SARS-CoV-2 Omicron subvariants (BA.1/2/3/4) were mapped (Fig.1a). 31 mutations from BA.1/2/3/4 in total can be revealed in our structures (Fig 1b). Among 20 mutations shared by four Omicron subvariants, 17 of them are clearly built. Five mutations (S371L, G496S, T547K, N856K, L981F) that only appeared in BA.1 are mapped except ins214EPE. All mutations shared by BA.2/3/4 (S371F, D405N), mutations shared by BA.1/2/3(Q493R), and all unique mutations from BA.4 (L452R, F486V) can be seen on the structure. As the common mutations shared by BA.1/3 are mostly located on NTD domain, only one of eight mutations were mapped (G446S). BA.2 and BA.4 have the highest number of shared mutations, and 3 of 8 were mapped (A27S, T376A, R408S). The other mutations are invisible due to local flexibility, such as N679K and P681H near the furin cleavage site (residues 682-685), and A67V, H69del, and V70del on the loops in the NTD.

### Distinct conformations of the S proteins from the four subvariants

Despite identical sample preparation and data processing procedures, the four complexes exhibit distinct conformational preferences (Fig. 2). In the 3D EM reconstruction for BA.1-SA, two RBDs are in the “up” conformation, while the third one is “down”. Consistent with previous studies, only the up RBDs are bound to ACE2-PD (*9*). In contrast, all three RBDs are “up” in BA.2-SA and BA.4-SA. Therefore, three ACE2-PD molecules are bound to the trimeric BA.2/4 S protein. BA.3-SA represents a mixture, with 81.7% selected particles in the all “up” states and the rest in the two up and one down conformation (Fig. 2).

**Fig. 2:**
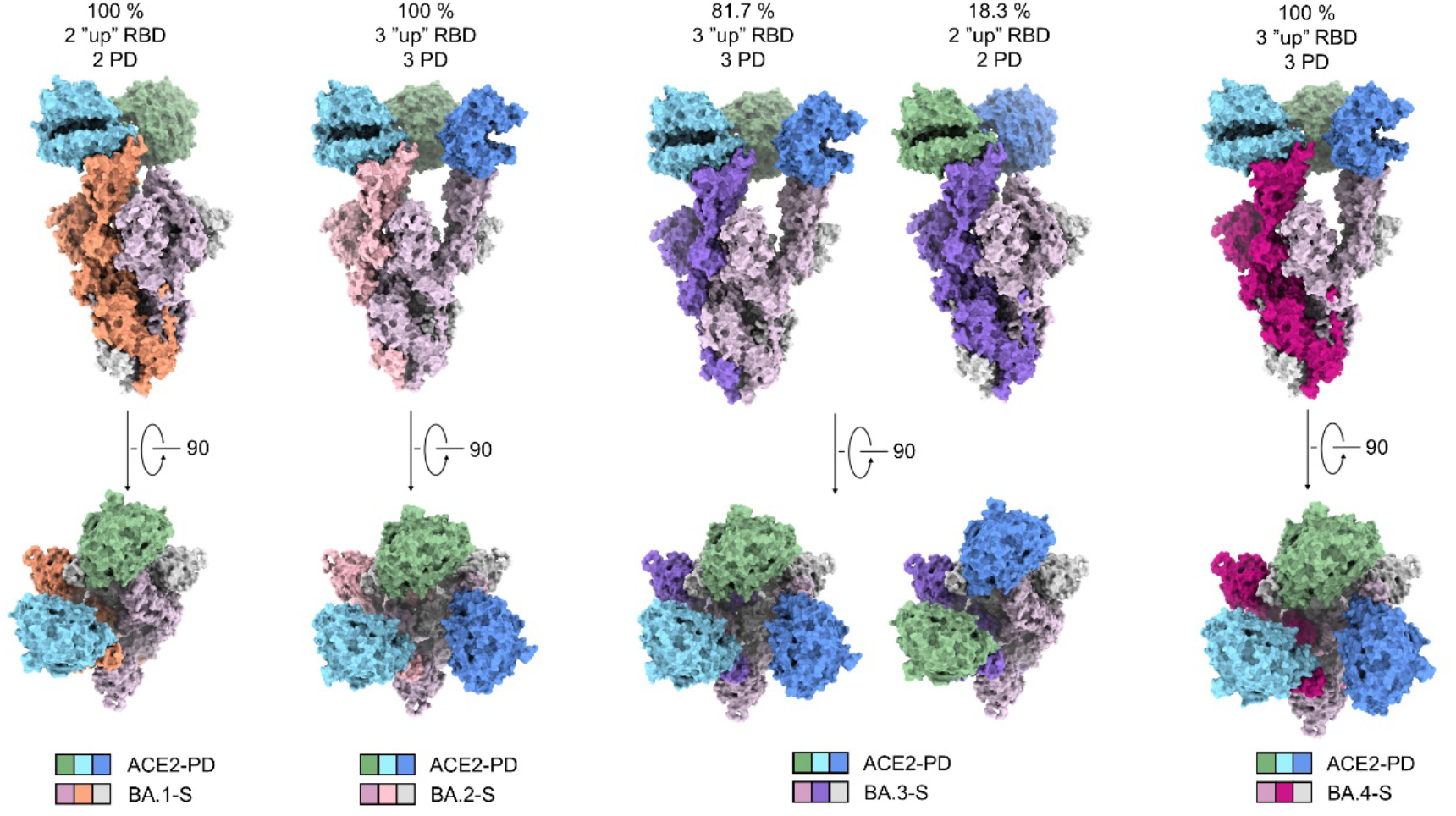
The trimeric S proteins from the four Omicron subvariants display distinct conformations and stoichiometric ratios with ACE2-PD. Shown here is surface presentation of domain-colored cryo-EM structures of S-ECD from Omicron BA.1, BA.2, BA.3 and BA.4 respectively in complex with the PD of ACE2. The three RBDs in BA.2-S and BA.4-S are all up and associate with three ACE2-PD. Two RBDs are up and one down in BA.1-S, and only the two up RBDs are associated with ACE2-PD. The majority (81.7%) of BA.3-S is identical with BA.2/4-S and the rest is the same as BA.1-S.

Of note, previous structures of WT S protein in complex with ACE2 show that WT-S tends to have either only one up and two “down” RBDs, or two “up” RBDs, among which only one is bound with ACE2-PD (67.7%) (*13, 21, 37*). The increasing tendency of more “up” RBDs from WT to Omicron sub-lineages is consistent with the enhanced infectivity of the Omicron subvariants, particularly BA.2 and BA.4. As BA.3-SA represents a mixture of the BA.1-SA and BA.2/4-SA conformations, following we will mainly focus on BA.2-SA and BA.4-SA for analysis.

### Mutations of BA.2 and BA.4 destabilize the down conformation of RBD

The molecular determinant for the “up” and “down” conformations of RBD is critical to understanding the altered infectivities of the Omicron subvariants. We compared the structures of the free S proteins from WT, BA.1, and BA.2, which are in the all three down state (*38-40*). To facilitate structural illustration, the RBDs in the neighboring protomers will be referred to as RBD and RBD’ (Fig. 3, top left).

**Fig. 3:**
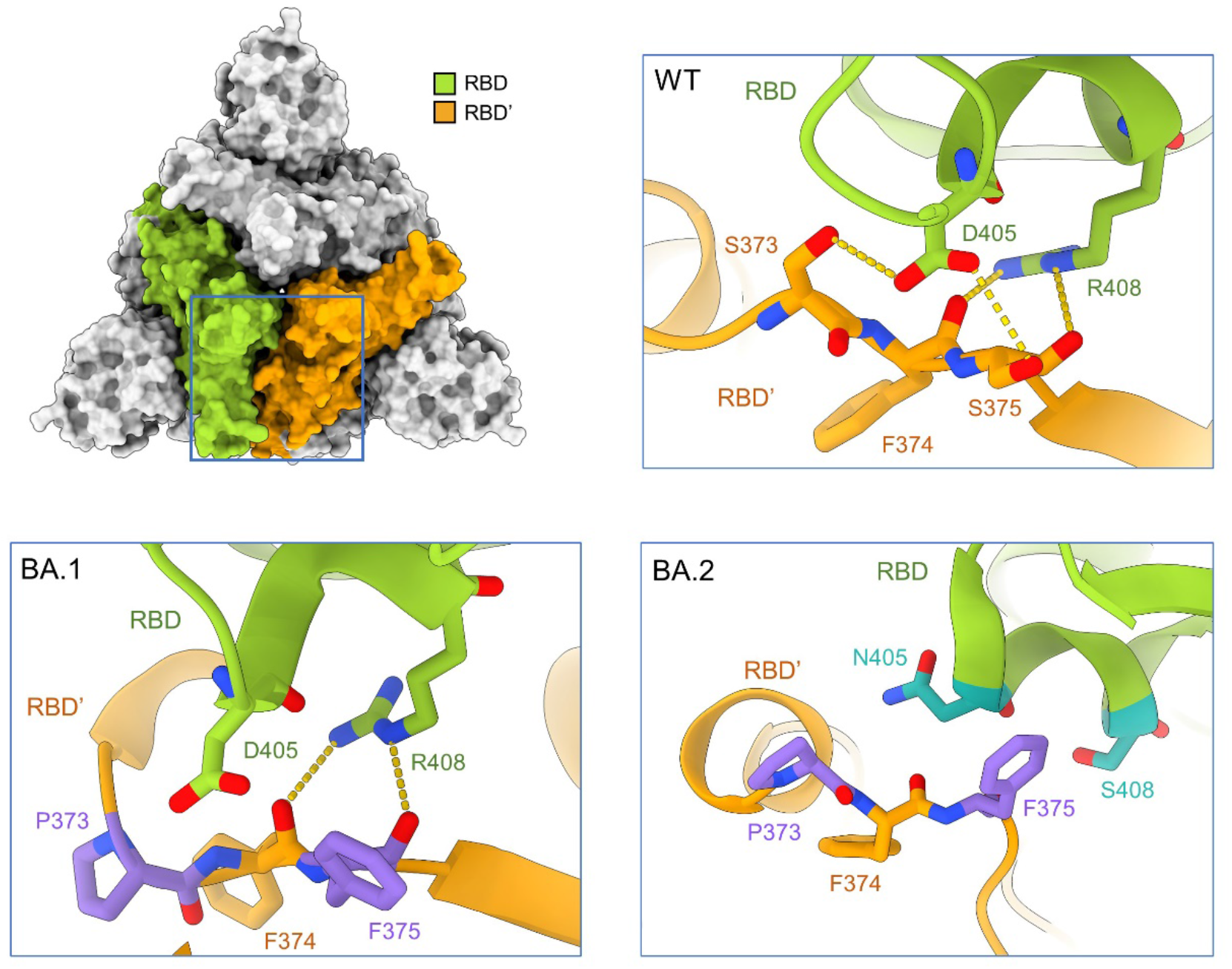
Mutations on the RBD-RBD’ interface in the Omicron subvariants may destabilize the down conformation, thus contributing to their enhanced infectivity by favoring the up conformation. Structure of the top view of the “all down” S-ECD from the original SARS-CoV-2 strain (designated as WT, PDB: 6ZB5) is shown on the top left. *Insets*: Enlarged views of the interface between the adjacent RBDs, labelled as RBD and RBD’, from WT (top right), BA.1 (bottom left, PDB: 7TF8), and BA.2 (bottom right, PDB: 7UB0). Residues on the interface are shown as sticks, green (RBD) or orange (RBD’) for invariant ones and dark green (RBD) or purple (RBD’) for mutated ones

In the WT-S protein, two hydrogen bonds (H-bonds) are formed between Asp405 and Arg408 in RBD and Ser373 and the backbone carbonyl oxygen groups of Phe374 and Ser375 in RBD’ (Fig. 3, top right). Mutations S373P and S375F in BA.1-S, which generate a more hydrophobic local environment, disrupt the interaction with Asp405 in the neighboring protomer. The carbonyl oxygen groups of Phe374 and Phe375 can still interact with the guanidinium group of Arg408 (Fig. 3, bottom left). In BA.2-S, however, the mutation R408S leads to the disruption of the interaction with the backbone carbonyl groups. In addition, D405N is less favored by the nearby amide groups in the neighboring protomer (Fig. 3, bottom right). The mutations, D405N and R408S, of BA.4 are same as BA.2, which might explain the same three “up” RBD conformation of structure in BA.4.

These mutations together weaken the packing between the neighboring RBDs in the down conformation, thus promoting the tendency of “up” conformation of the S protein and suitable for ACE2 binding. The further disrupted RBD-RBD’ interface in the down conformation thus affords a clue to the all “up” conformation observed in our structure (Fig. 2).

### Several Omicron mutations strengthen the interaction with ACE2

While the different conformational preferences of the trimeric S protein may account for the increased infectivity of the Omicron subvariant, particularly BA.2 and BA.4, they cannot explain the higher affinity between the RBD of all four Omicron subvariants and ACE2-PD, as the measurement was performed with isolated monomeric domain (Supplementary Fig. 1). We thereby scrutinized the mutations on the interface with ACE2.

Structural analysis has identified 9 RBD residues on the interface with ACE2. Sequence alignment of the RBD from WT, Delta, and Omicron subvariants reveals that only 3 residues, Tyr453, Gln474, and Thr500, remain identical in all the variants, while 5 mutated in BA.1, BA.2, and BA.3 subvariants, including K417N, S477N, Q493R, Q498R, and N501Y. BA.1 contains an additional mutation G496S and BA.4 contains an additional mutation F486V and keep the Gln493 as WT S protein, that might affect the interaction net. These mutations remodel the interaction network between RBD and ACE2 (Fig. 4, Supplementary Fig. 8).

**Fig. 4:**
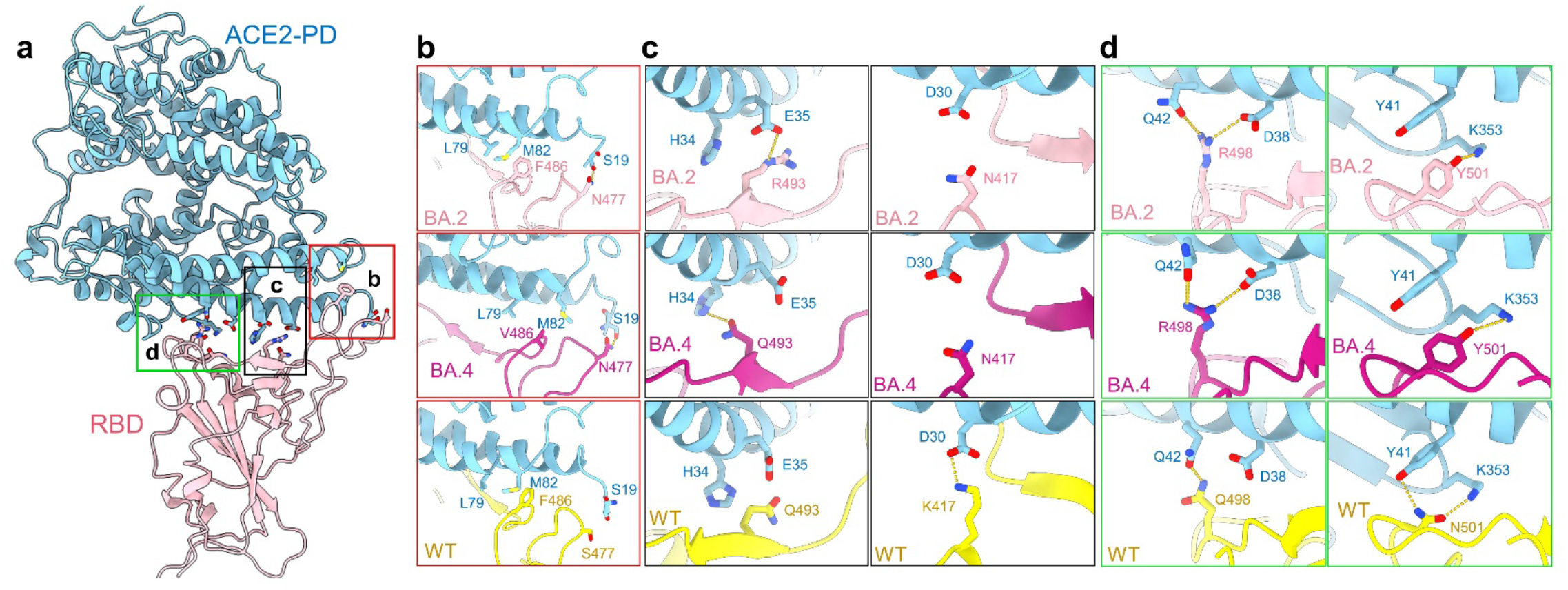
Mutations on the interface of ACE2 and BA.2/4-S underlie their enhanced affinity. **a**, Point mutations of BA.2-S strengthen its interaction with ACE2. Detailed comparison of the ACE2-interacting residues between BA.2, BA.4 and WT S protein are shown in panels **b-d. b**, S477N of BA.2/4-S results in an additional hydrogen bond with Ser19 of ACE2. F486V keeps the Waals forces with Met82 and Leu79 of ACE2 **c**, Q493R of BA.2-S results in a salt bridge with ACE2-Glu35 and Gln493 of BA.4 interacts with ACE2-His34, although K417N weakens the original interaction with ACE2-Asp30. **d**, Q498R of BA.2/4 results in a salt bridge with ACE2-Asp38, and N501Y leads to a rearrangement of the local interactions with Tyr41 and Ly353 in ACE2. The PDB ID for the WT structure is 6M17.

Comparing with WT, mutations S477N, Q493R, and Q498R in Omicron RBD lead to new polar interactions with Ser19, Glu35, and Asp38 in ACE2, respectively. These additional contacts, particularly the strong salt bridges mediated by Arg493 and Arg498, may not only compensate for the lost interactions of Lys417 and Asn501 in the WT S protein respectively with Asp30 and Tyr41 in ACE2, but result in a 2-4 fold net increase of the affinity (Fig.4b-d). Besides, mutation F486V of BA.4 remains the hydrophobic interaction with Leu79 and Met82 of ACE2 and Gln493 is H-bonded with His34 of ACE2, which might be affected by the local environment of L452R mutation in BA.4 (Fig.4b,4c).

### The immune evasion mechanism of S309 and S2K146

Next we examined the resolved mutations to look for clues to the immune evasion mechanism of Omicron subvariants. Many of the surface-mapping mutations (Fig. 1b) have been analyzed previously (*13, 41*). To avoid redundancy, here we mainly focus on the distinct responses to antibody S309, which can still neutralize BA.1, but not the BA.2 and BA.4 subvariants, and antibody S2K146, which could still neutralize BA.1, BA.2, and BA.3, but not the BA.4 (*26, 30, 31*).

The authorized monoclonal antibody S309 has been shown to effectively neutralize several SARS-CoV-2 variants, including BA.1. However, its neutralizing activity against BA.2 and BA.4 is largely compromised (*23*). In contrast, another broadly neutralizing antibody, LY-CoV1404, is still potent in neutralizing all Omicron sub-lineages (*23, 31, 32*). Both S309 and LY-CoV1404 belong to Class 3 that target outside ACE2 binding site and recognize both “up” and “down” RBDs.

Among all mutations on the RBD, only G339D is positioned in the epitope of S309. However, it is shared by BA.1, BA.2, and BA.4 (Fig. 5a and Supplementary Fig. 8). Therefore, this mutation does not account for the different sensitivities of BA.1, BA.2, and BA.4 to S309. We therefore compared the maps for BA.1-SA, BA.2-SA and BA.4-SA for potential allosteric mutation sites. We observed an evident shift of the glycosylation moieties linked to Asn343 among these three maps. Furthermore, the conformation of this glycan is nearly identical between BA.1 and WT (Fig. 5b, Supplementary Fig. 7h). This difference can be potentially important as the Asn343 glycan is directly involved in S309 recognition (Fig. 5a)(*42*).

**Fig. 5:**
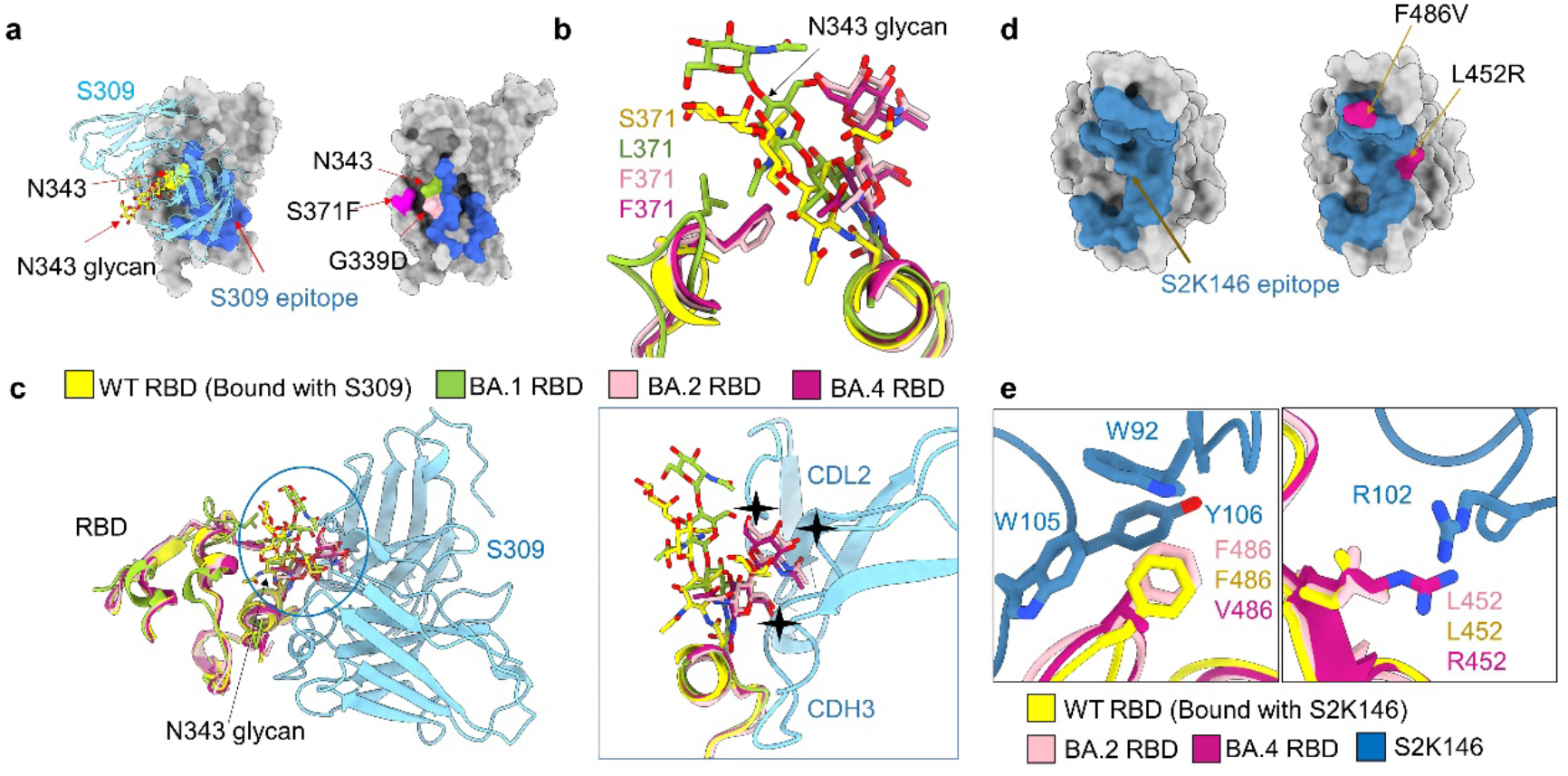
Potential mechanism for S309 and S2K146 escaping neutralization. **a**, There is no BA.2/4-unique mutation on the S309 epitope. *Left*: A glycan on Asn343, shown as yellow sticks, is recognized by antibody S309. Shown here is the structure of WT RBD bound to antibody S309 (PDB code: 6WPT). The S309 epitope is colored blue. *Right*: Omicron subvariant mutations in the vicinity of the S309 epitope. G339D, which is common to BA.1, BA.2 and BA.4, is the only one mapped to the S309 epitope. S371F is a mutation in BA.2, BA.3 and BA.4, but missing in BA.1. **b**, The glycosylation moieties of Asn343 of RBD is pushed away due to the steric hindrance as a result of S371F mutation in BA.2/4-S. **c**, Mutation S371F may underlie BA.2, BA.3 and BA.4’s evasion from S309 neutralization by altering the conformation of Asn343 glycan. Structures of the RBD from the WT in complex with antibody S309, BA.1, BA.2 and BA.4 are superimposed (left). The Asn343 glycan in BA.1 exhibits a similar conformation as that in WT, while that in BA.2 and BA.4 has to move away due to the potential steric clash with Phe371. The shifted Asn343 glycan in BA.2 and BA.4 would clash with antibody S309 (right), hence impeding its recognition of the epitope. **d**, There are two unique mutations of BA.4 on the S2K146 epitope. Left: The S2K146 epitope is colored steel blue (PDB code: 7TAS). Right: unique mutations of BA.4 in the vicinity of the S2K146 epitope. **e**, F486V in BA.4 disrupt the pie-pie interaction net that involves the Phe486 of RBD, Trp92 of light chain, Trp105 and Tyr106 of heavy chain and the L452R could cause a clash with Arg102 of heavy chain.

Scrutiny of the local environment suggests that the shift of the N343-glycan in BA.2-SA and BA.4-SA is a direct consequence of the mutation S371F, which is found in BA.2, BA.3, and BA.4, but not BA.1. The bulky hydrophobic ring of Phe371 presents a steric hindrance to the polar sugar moieties nearby. Therefore, the N343-glycan has to move away. Notably, this shift would result in a clash with the antibody S309, making the binding less favored (Fig. 5c, Supplementary Fig. 7h). Our comparative structural study thus reveals an allosteric mechanism of the escape from S309 by BA.2, BA.3, and BA.4 through the single point mutation S371F, which is not directly positioned on the S309 epitope.

Antibody S2K146 can neutralize BA.2 and BA.3, but significantly lose its neutralizing activity against BA.4. The compromised neutralization of S2K146 to BA.4 might be directly affected by the mutations, L452R and F486V of BA.4 (Fig.5d). When aligned the epitopes of S2K146 with WT, BA.2, and BA.4, the F486V obviously disrupts the pie-pie interaction net that involves the Phe486 of RBD, Trp92 of light chain, Trp105 and Tyr106 of heavy chain (Fig.5e, left). The L452R could cause a clash with Arg102 of heavy chain (Fig.5e, right).

LY-CoV1404 preservs neutralizing activity for all five Omicron subvariants. It is noted that four Omicron mutations, N440K, G446S, Q498R, and N501Y, are mapped to the epitope of LY-CoV1404 (Supplementary Fig. 9a). Instead of disrupting interactions with the antibody, Lys440 and Arg498 of BA.2 form H-bonds with Tyr35 and Thr96 of LY-CoV1404, respectively. The mutation N501Y does not affect the interaction as neither Asn nor Tyr directly participates in the interaction with the antibody. Besides, G446S, a common mutation in BA.1 and BA.3, might lead to the interaction between Ser446 and Arg60 of heavy chain in LY-CoV1404 (Supplementary Fig. 9b). As seen, none of these mutations impair the interaction with the antibody, explaining the broad neutralizing mechanism of LY-CoV1404.

## Discussion

In this study, we attempt to provide clues to two critical questions regarding the Omicron variants through comparative structural analysis of ACE2-bound S proteins from BA.1/2/3/4. First, the molecular basis for the increased transmissibility of Omicron subvariants, particularly BA.2 and BA.4. Second, the mechanism for distinct responses to antibody neutralization.

In addition to the varied affinity with the cellular receptor ACE2, the switch to control the “up” and “down” states of RBD is critical for the infectivity of SARS-CoV-2. Our structural investigation reveals that in the presence of the PD of ACE2, RBDs in all three protomers of BA.2/4 S protein exhibit the “up” conformation, whereas only two are “up” in BA.1-S. This observation may partially account for the increased infectivity of BA.2 and BA.4. Previous studies have shown that cleavage of the furin cleavage site promotes the “up” conformation of RBD (*21, 43*). Here we discover that a preference for the “up” state of RBD bound with ACE2 may be achieved through destabilizing the down state packing and facilitating ACE2 binding. Therefore, the Omicron subvariants may acquire enhanced infectivity through both tighter binding to ACE2 and increased tendency for the “up” state. Detailed analysis of the disruptive mutations on the RBD interface in their “down” state is consistent with their differential preference for the “up” state.

The COVID-19 pandemic has continued to spread across the world for over two years (*44*). Under the selective pressure of the immune system, variants of SARS-CoV-2 have evolved to evade the antibody immunity elicited by the vaccine immunization or natural viral infection (*45-49*). A large number of mutations of BA.1,vBA.2, BA.3, and BA.4 are distributed on the hotspots of epitopes, providing a mechanistic basis for the immune evasion from many therapeutic antibodies and vaccinations (Fig. 1b). Out of expectation, an allosteric mechanism is discovered to allow BA.2, BA.3, and BA.4 to escape the neutralization of S309 through the mutation of S371F, which alters the conformation of a nearby glycan to hinder antibody binding. This discovery highlights the role of glycosylation in antibody recognition. Besides, the additional mutations of BA.4 further reduce the neutralization activity of BA.4, suggesting an unexpected fitness of SARS-CoV-2.

Taken together, our studies reveal the molecular basis for the recognition of the S proteins of Omicron subvariants by the ACE2 receptor. Our analyses presented here provide molecular clues for the higher infectivity and antibody evasion of Omicron sub-lineages and shed light on the therapeutic interventions against SARS-CoV-2 variants.

## Acknowledgments

We thank the Cryo-EM Facility and Supercomputer Center of Westlake University for providing cryo-EM and computation support, respectively. This work was funded by the start-up funds from the Southern University of Science and Technology (To R.Y.).

## Author contributions

R.Y. conceived the project. R.Y., Y.S., and Y.L. designed the experiments. All authors did the experiments. R.Y., Y.Z., Y.S., and Y.L. contributed to data analysis. R.Y. wrote the manuscript.

## Data and materials availability

Atomic coordinates and cryo-EM density maps of Omicron BA.2 S protein in complex with PD of ACE2 (PDB: 7Y1Y, whole map: EMD-33575, map focused on the interface between BA.2 RBD and ACE2: EMD-33579). Two “up” RBD state of BA.3 S protein in complex with PD of ACE2 (PDB: 7Y20, whole map: EMD-33577) and three “up” RBD state of BA.3 S protein in complex with PD of ACE2 (PDB: 7Y1Z, whole map: EMD-33576, map focused on the interface between BA.3 RBD and ACE2: EMD-33580) have been deposited to the Protein Data Bank (http://www.rcsb.org) and the Electron Microscopy Data Bank (https://www.ebi.ac.uk/pdbe/emdb/), respectively. BA.4 S protein in complex with PD of ACE2 (PDB: 7Y21, whole map: EMD-33578, map focused on the interface between BA.4 RBD and ACE2: EMD-33581). The other PDB and EMDB IDs can be found in Supplementary information, Supplementary Table 1.

## Supplementary Materials

Materials and Methods; Supplementary Figures S1-S9; Supplementary Table 1.

