## supplemental information for "Structural basis for the enhanced infectivity and immune evasion of Omicron subvariants"

### Materials and methods

#### Protein expression and purification

The extracellular domain (ECD) (1-1208 a.a) of S protein of SARS-CoV-2 Omicron variant BA.1/2/3/4 were cloned into the pCAG vector (Invitrogen) with six proline substitutions at residues 817, 892, 899, 942, 986 and 987 and a C-terminal T4 fibritin trimerization motif followed by 10×His tag, respectively. A "GSAS" mutation at residues 682 to 685 was introduced into ECD to prevent the host furin protease digestion. These constructs were hereafter referred to as BA.1/2/3/4-S.

The receptor binding domain (RBD) (319-541 a.a) of S protein from SARS-CoV-2 WT strain and Omicron variant BA.1/2/3/4 were cloned into the pCAG vector (Invitrogen) with an N-terminal signal peptide of secreted luciferase and a C-terminal 6×His tag, respectively. These residue numbers mentioned above are that relative to the spike (WT). The peptidase domain (PD) (19-615 a.a) of human ACE2 was also cloned into the pCAG vector (Invitrogen) with an N-terminal signal peptide of secreted luciferase and a C-terminal Flag tag. The mutants were generated with a standard two-step PCR-based strategy. All the plasmids used to transfect cells were prepared by GoldHi EndoFree Plasmid Maxi Kit (CWBIO).

The recombinant protein was overexpressed using the HEK293F mammalian cells at 37°C under 5% $CO_2$ in a Multitron-Pro shaker (Infors, 130 rpm). When the cell density reached $2.0 \times 10^6$ cells/mL, the plasmid was transiently transfected into the cells. To transfect one liter of cell culture, about 1.5 mg of the plasmid was premixed with 3 mg of polyethylenimines (PEIs) (Polysciences) in 50 mL of fresh medium for 15 mins before adding to cell culture. Cells were removed and medium was collected by centrifugation at 4000×g for 15 mins after sixty hours transfection.

The secreted ECD and RBD of S protein were purified by Ni-NTA affinity resin (Qiagen). The nickel resin loaded was rinsed with the wash buffer 1 containing 25 mM HEPES (pH 7.0), 500 mM NaCl and washed with wash buffer 2 containing 25 mM HEPES (pH 7.0), 150 mM NaCl and 30 mM imidazole. Protein was eluted by wash buffer 2 plus 270 mM imidazole. Then the Ni-NTA eluent of ECD was subjected to size-exclusion chromatography (Superose 6 Increase 10/300 GL, GE

Healthcare) in buffer containing 25 mM HEPES (pH 7.0), 150 mM NaCl. The peak fractions were collected and stored at -80°C. The Ni-NTA eluent of RBD was subjected to size-exclusion chromatography (Superose 6 Increase 10/300 GL, GE Healthcare) in PBS buffer (pH 7.4, gibco) with 0.04% Tween-20. The factions were collected for measurement of RBD binding to human ACE2-PD by biolayer interferometry.

The secreted PD was purified by anti-FLAG M2 affinity resin (Sigma Aldrich). After loading two times, the anti-FLAG M2 resin was washed with the wash buffer 3 containing 25 mM HEPES (pH 7.0), 150 mM NaCl. The protein was eluted with the wash buffer 3 plus 0.2 mg/mL flag peptide. The eluent of PD was then concentrated and subjected to size-exclusion chromatography (Superdex 200 Increase 10/300 GL, GE Healthcare) in buffer containing 25 mM HEPES (pH 7.0), 150 mM NaCl.

The BA.1/2/3/4-S was incubated with PD at a molar ratio of about 1:6 for one hour. To remove excessive PD, the mixture was subjected to size-exclusion chromatography (Superose 6 Increase 10/300 GL, GE Healthcare) in buffer containing 25 mM HEPES (pH 7.0), 150 mM NaCl. The peak fractions containing protein complex were collected for EM analysis.

**Measurement of Omicron subvariants and WT strain RBD binding to human ACE2-PD by biolayer interferometry**

The binding between PD and RBD of Omicron subvariants were performed using Octet Red96e (ForteBio), and the RBD of WT strain was measured as well at the same time. The PD of ACE2 was biotinylated using biotinylation kit (Genemore, 1828M) and loaded to octet SA biosensor (Sartorius). The association and dissociation of PD-coated biosensors with different concentrations of RBD of SARS-CoV-2 S protein were recorded in binding buffer (PBS pH 7.4, 0.04% Tween-20). Data was analyzed by Octet Data Analysis HT 12.0 software. Reference sample and reference sensor were subtracted, and K D values were analyzed using a 1:1 global fit model. Data were plotted using Prism V8.0 software (GraphPad).

**Cryo-EM sample preparation and data acquisition**

The BA.1/2/3/4-SA were concentrated to ~1.5 mg/mL and applied to the grids. Aliquots (3.3 μL) of the protein were placed on glow-discharged holey carbon grids (Quantifoil Au R1.2/1.3). The grids were blotted for 3.0 s or 3.5 s and flash-frozen in liquid ethane cooled by liquid nitrogen with Vitrobot (Mark IV, Thermo Fisher Scientific). The prepared grids were transferred to a Titan Krios operating at 300 kV equipped with Gatan K3 detector and GIF Quantum energy filter. Movie stacks were automatically collected using AutoEMation[1], with a slit width of 20 eV on the energy filter and a defocus range from -1.4 μm to -1.8 μm in super-resolution mode at a nominal magnification of 81,000×. Each stack was exposed for 2.56 s with an exposure time of 0.08 s per frame, resulting in a total of 32 frames per stack. The total dose rate was approximately 50 $e^-$/$Å^2$ for each stack. The stacks were motion corrected with MotionCor2[ref.2] and binned 2-fold, resulting in a pixel size of 1.087 Å/pixel. Meanwhile, dose weighting was performed[3]. The defocus values were estimated with Gctf[4].

**Data processing**

The Cryo-EM structure of S protein from BA.1-SA has been solved firstly[5] and identical protocol was applied to the complex of BA.2, BA.3 and BA.4. Particles for S-ECD bound with PD of ACE2 were automatically picked using Relion 3.0.6[ref.6-9] from manually selected micrographs. After 2D classification with Relion, good particles were selected and subject to multiple cycle of heterogeneous refinement without symmetry using cryoSPARC[10]. The good particles were selected and subjected to Local CTF Refinement with C1 symmetry, Non-uniform Refinement, resulting in the 3D reconstruction for the whole structures. For interface between RBD of S-ECD and PD, the dataset was subject to focused refinement with adapted mask and reference on RBD-PD sub-complex to improve the map quality. Then the dataset of four RBD-PD sub-complexes were combined and subject to focused refinement with Relion and then were subject to Local Refinement with cryoSPARC, resulting in the 3D reconstruction of better quality on the interface between S-ECD and PD.

The resolution was estimated with the gold-standard Fourier shell correlation 0.143 criterion[11] with high-resolution noise substitution[12]. Refer to Materials and Methods, Extended Data Figs.2-7 and Extended Data Table 1 for details of data collection and processing.

**Model building and structure refinement**

For model building of the complex of BA.2/3/4-SA, the atomic model of the T-ACE-S(p) (PDB ID: 7CT5) were used as templates, which were molecular dynamics flexible fitted[13] into the whole cryo-EM map of the complex and the focused-refined cryo-EM map of the RBD-PD sub-complex, respectively. Each residue was manually checked with the chemical properties taken into consideration during model building. Several segments, whose corresponding densities were invisible, were not modeled. Structural refinement was performed in Phenix[14] with secondary structure and geometry restraints to prevent overfitting. To monitor the potential overfitting, the model was refined against one of the two independent half maps from the gold-standard 3D refinement approach. Then, the refined model was tested against the other map. Statistics associated with data collection, 3D reconstruction and model building were summarized in Table S1.

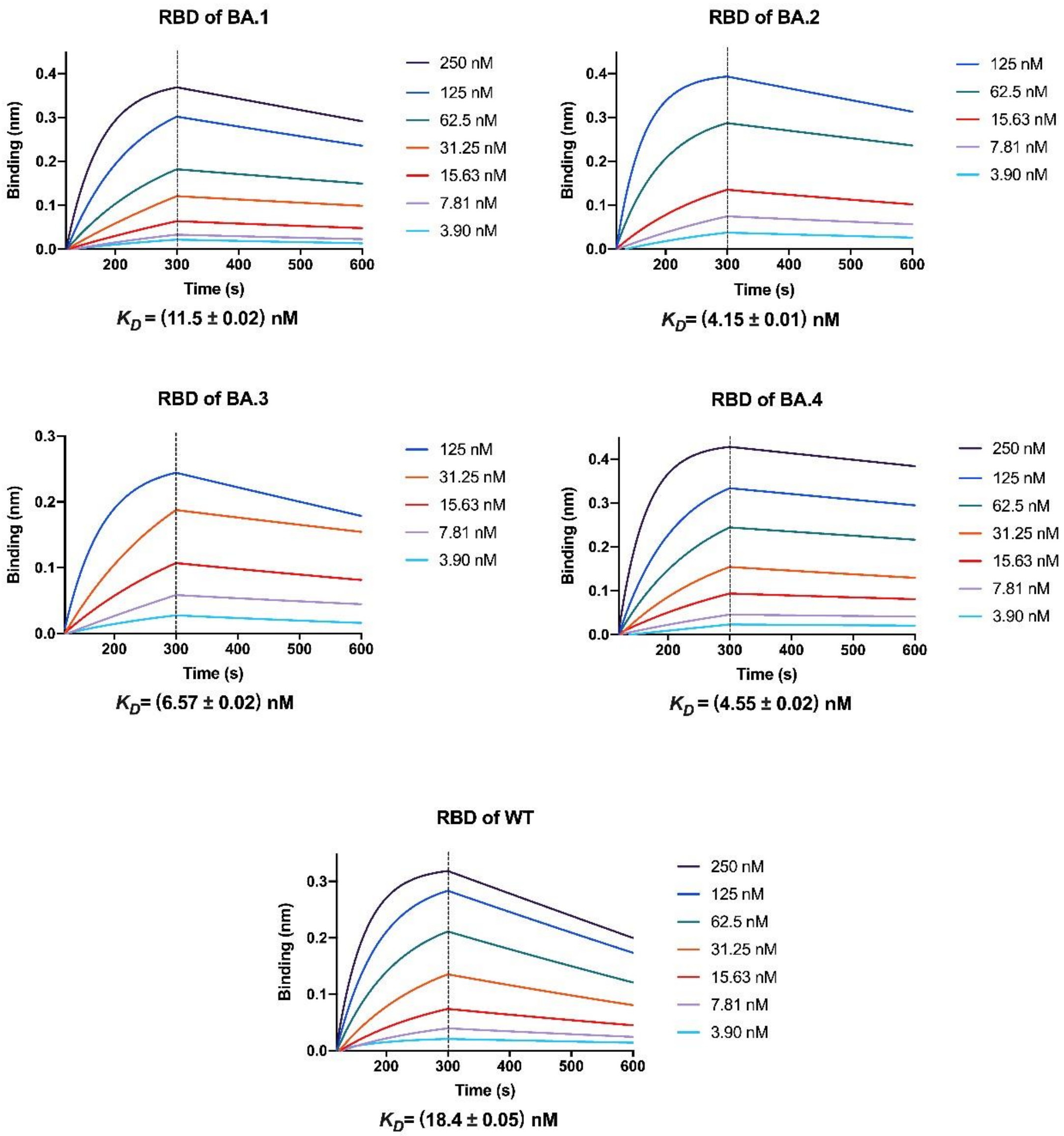


**Supplementary Fig.1**

**The RBD of BA.2/4-S has slightly higher affinity with the host receptor ACE2.**

Binding affinities of the S-RBD from the four Omicron subvariants with the peptidase domain of ACE2 (ACE2-PD). The association and dissociation of S-RBD, applied at different concentrations, with ACE2 PD-coated Streptavidin biosensors was measured using Bio-Layer Interferometry (BLI). $K_D$ was analyzed with Octet Data Analysis HT 12.0 software.

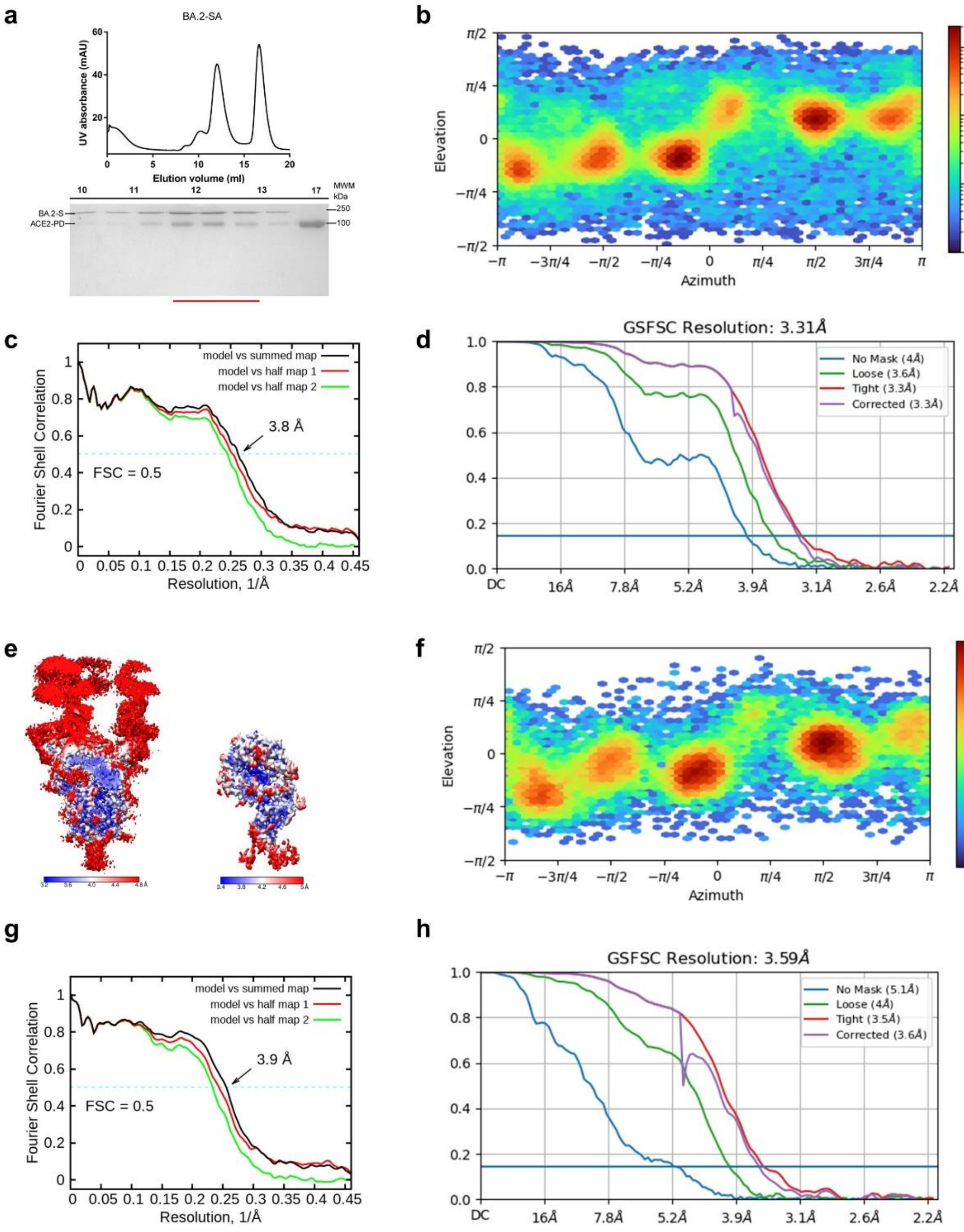


**Supplementary Fig.2**

**Cryo-EM analysis of S-ECD from BA.2-SA.**

**a,** Representative SEC purification of the BA.2-SA. SDS–PAGE was visualized by Coomassie blue staining and fractions for cryo-EM analysis were marked by red line. **b,** Euler angle distribution in the final 3D reconstruction of overall map. **c,** FSC curve of the refined model of BA.2-SA versus the overall structure that it is refined against (black); of the model refined against the first half map versus the same map (red); and of the model refined against the first half map versus the second half map (green). The small difference between the red and green curves indicates that the refinement of the atomic coordinates did not suffer from overfitting. **d,** FSC curve of BA.2-SA. **e,** Local resolution map for the 3D reconstruction of the overall structure and the structure of RBD-PD. **f,** Euler angle distribution in the final 3D reconstruction of RBD-PD sub-complex. **g,** FSC curve of the refined model of RBD-PD sub-complex versus the overall structure that it is refined against (black); of the model refined against the first half map versus the same map (red); and of the model refined against the first half map versus the second half map (green). The small difference between the red and green curves indicates that the refinement of the atomic coordinates did not suffer from overfitting. **h,** FSC curve of RBD-PD sub-complex.

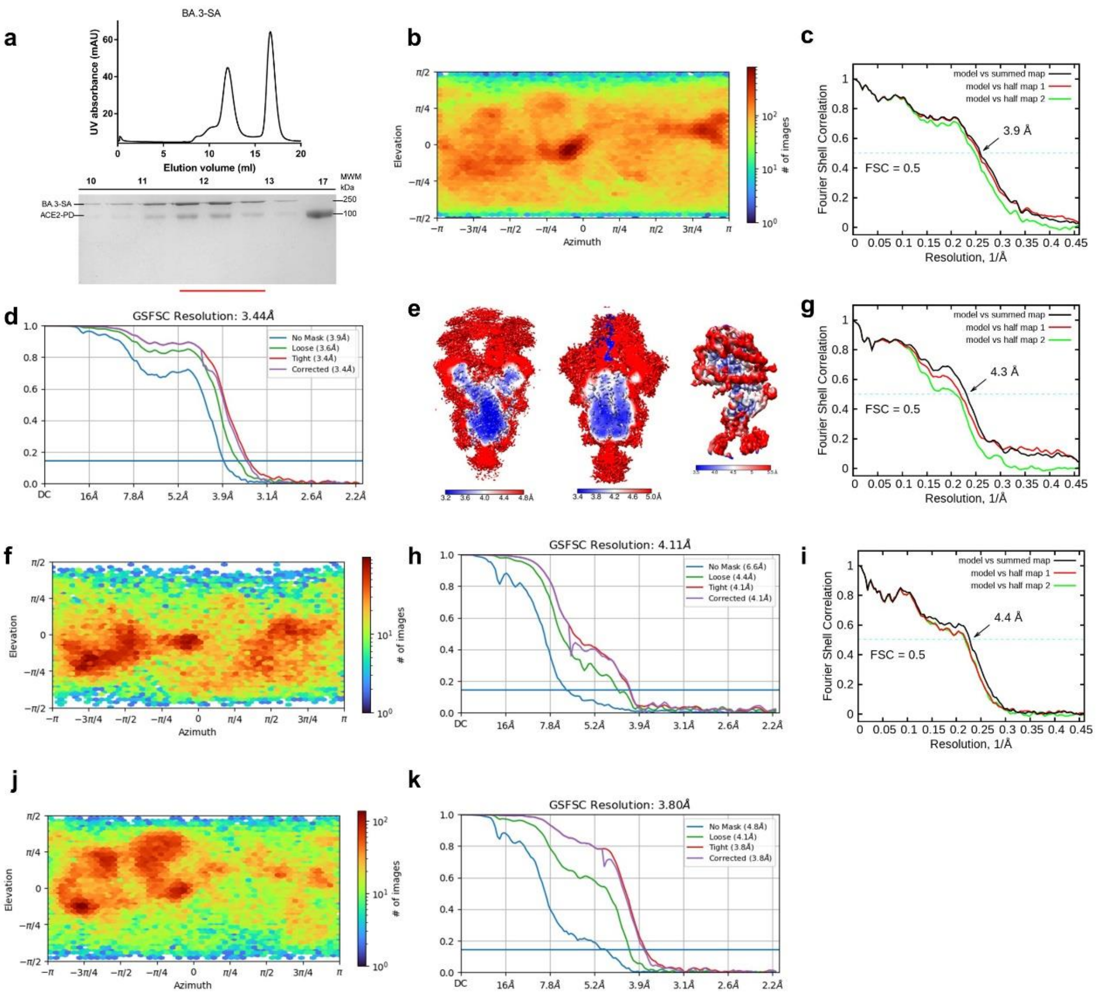


**Supplementary Fig.3**

**Cryo-EM analysis of BA.3-SA.**

**a,** Representative SEC purification of the BA.3-SA. SDS–PAGE was visualized by Coomassie blue staining and fractions for cryo-EM analysis were marked by red line. **b,** Euler angle distribution in the final 3D reconstruction of overall map of 3 "up" RBD BA.3-S in complex with 3 PD. **c,** 3 "up" RBD FSC curve of the refined model of BA.3-S in complex with 3 PD versus the overall structure that it is refined against (black); of the model refined against the first half map versus the same map (red); and of the model refined against the first half map versus the second half map (green). The small difference between the red and green curves indicates that the refinement of the atomic coordinates did not suffer from overfitting. **d,** FSC curve of 3 "up" BA.3-S in complex with 3 PD. **e,** Local resolution map for the 3D reconstruction of BA.3-S in complex with PD at 3 "up" RBD 3 PD state (left), 2 "up" RBD 2 PD state (middle) overall structure and RBD-PD sub-complex (right). **f,** Euler angle distribution in the final 3D reconstruction of RBD-PD sub-complex. **g,** FSC curve of the refined model of RBD-PD sub-complex versus the overall structure that it is refined against (black); of the model refined against the first half map versus the same map (red); and of the model refined against the first half map versus the second half map (green). The small difference between the red and green curves indicates that the refinement of the atomic coordinates did not suffer from overfitting. **h,** FSC curve of RBD-PD sub-complex. **(i)** Euler angle distribution in the final 3D reconstruction of overall map of 2 “up” RBD BA.3-S in complex with 2 PD. **j,** 2 “up” RBD FSC curve of the refined model of BA.3-S in complex with 2 PD versus the overall structure that it is refined against (black); of the model refined against the first half map versus the same map (red); and of the model refined against the first half map versus the second half map (green). The small difference between the red and green curves indicates that the refinement of the atomic coordinates did not suffer from overfitting. **k,** FSC curve of 2 “up” RBD BA.3-S in complex with 2 PD.

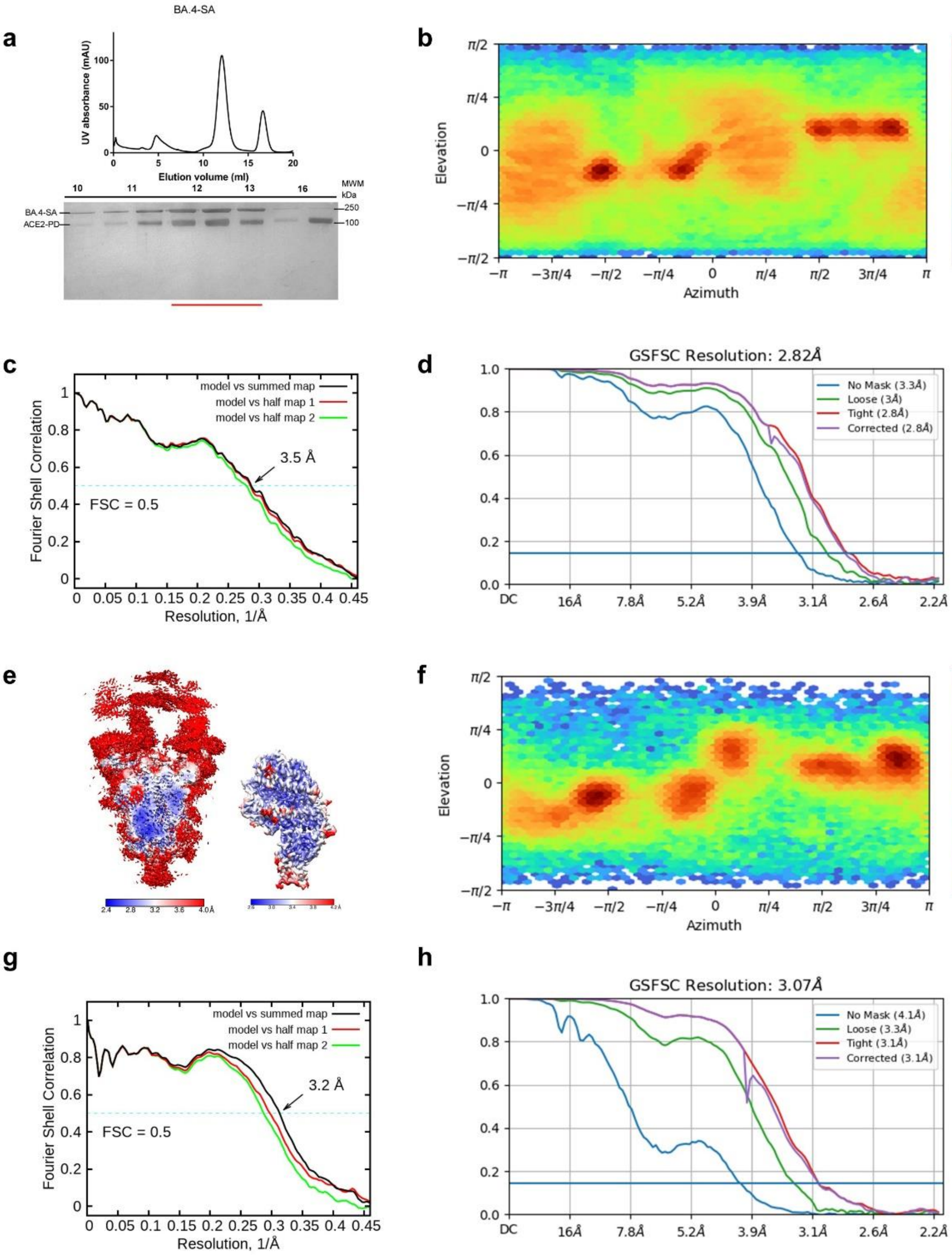

**Supplementary Fig.4**

**Cryo-EM analysis of S-ECD from BA.4-SA.**

**a,** Representative SEC purification of the BA.4-SA. SDS–PAGE was visualized by Coomassie blue staining and fractions for cryo-EM analysis were marked by red line. **b,** Euler angle distribution in the final 3D reconstruction of overall map. **c,** FSC curve of the refined model of BA.4-SA versus the overall structure that it is refined against (black); of the model refined against the first half map versus the same map (red); and of the model refined against the first half map versus the second half map (green). The small difference between the red and green curves indicates that the refinement of the atomic coordinates did not suffer from overfitting. **d,** FSC curve of BA.4-SA. **e,** Local resolution map for the 3D reconstruction of the overall structure and the structure of RBD-PD. **f,** Euler angle distribution in the final 3D reconstruction of RBD-PD sub-complex. **g,** FSC curve of the refined model of RBD-PD sub-complex versus the overall structure that it is refined against (black); of the model refined against the first half map versus the same map (red); and of the model refined against the first half map versus the second half map (green). The small difference between the red and green curves indicates that the refinement of the atomic coordinates did not suffer from overfitting. **h,** FSC curve of RBD-PD sub-complex.

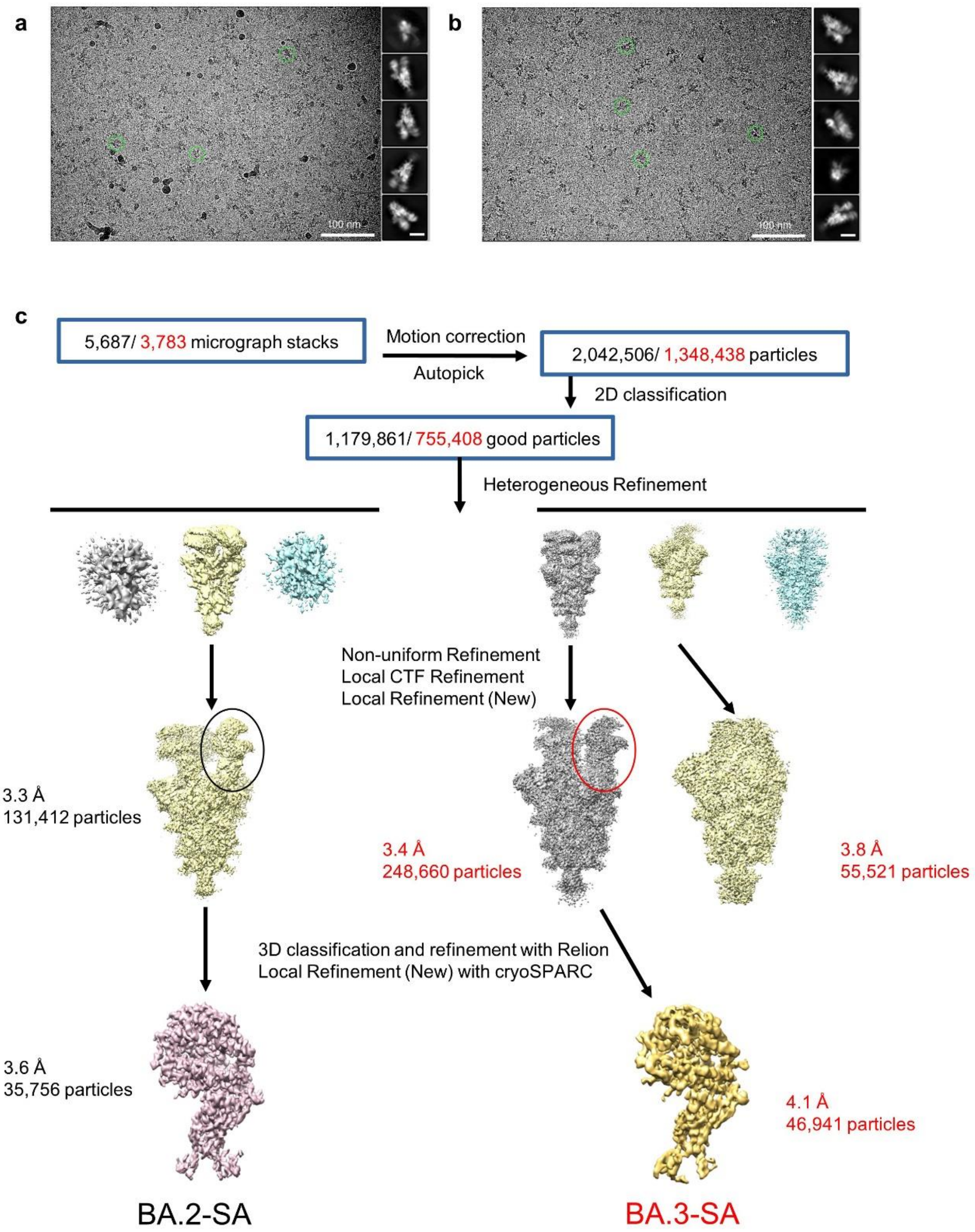


**Supplementary Fig.5**

**Flowchart of BA.2/3 for cryo-EM data processing.**

Please refer to the 'Data Processing' in Methods section for details.

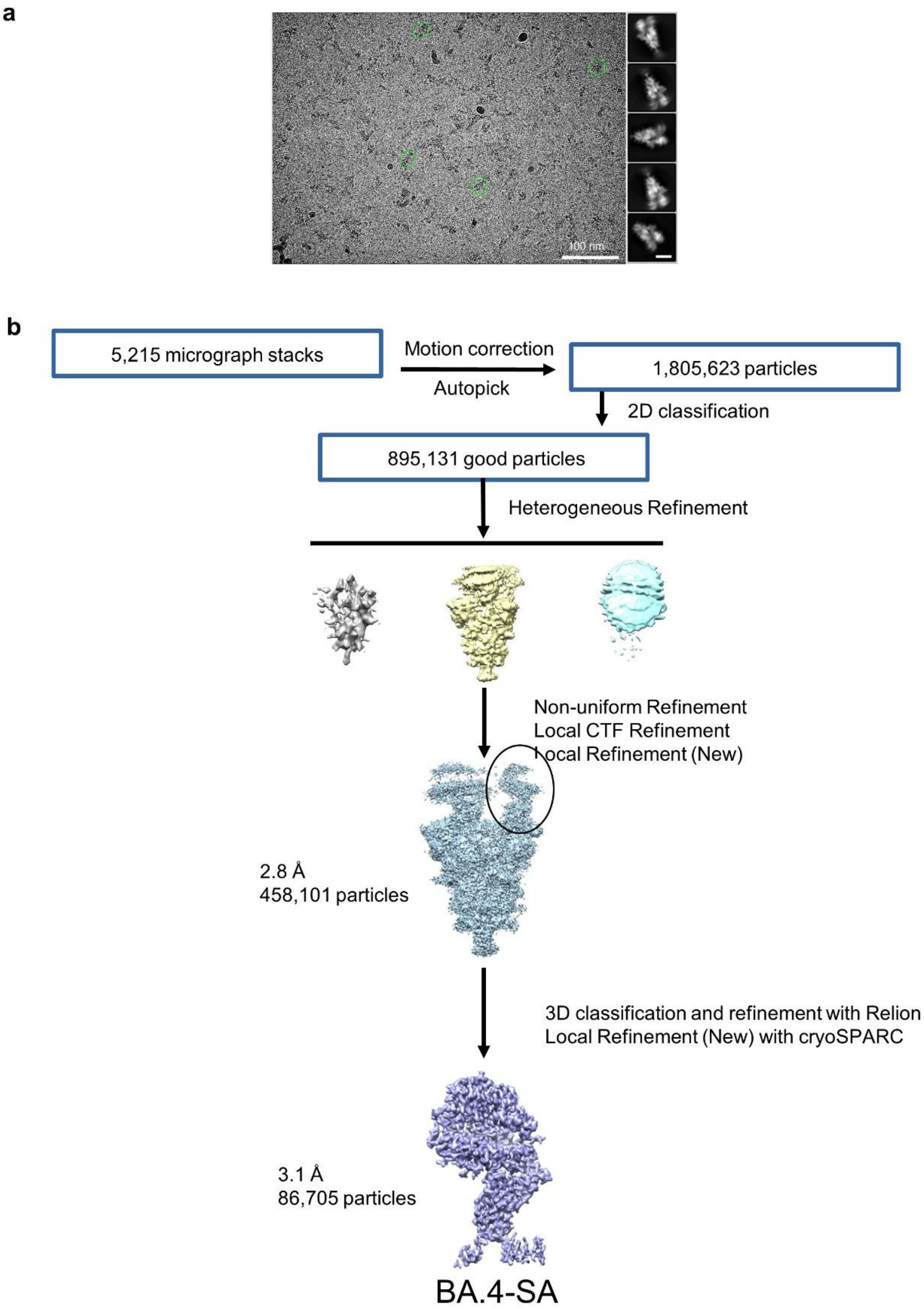


**Supplementary Fig.6**

**Flowchart of BA.4 for cryo-EM data processing.**

Please refer to the 'Data Processing' in Methods section for details.

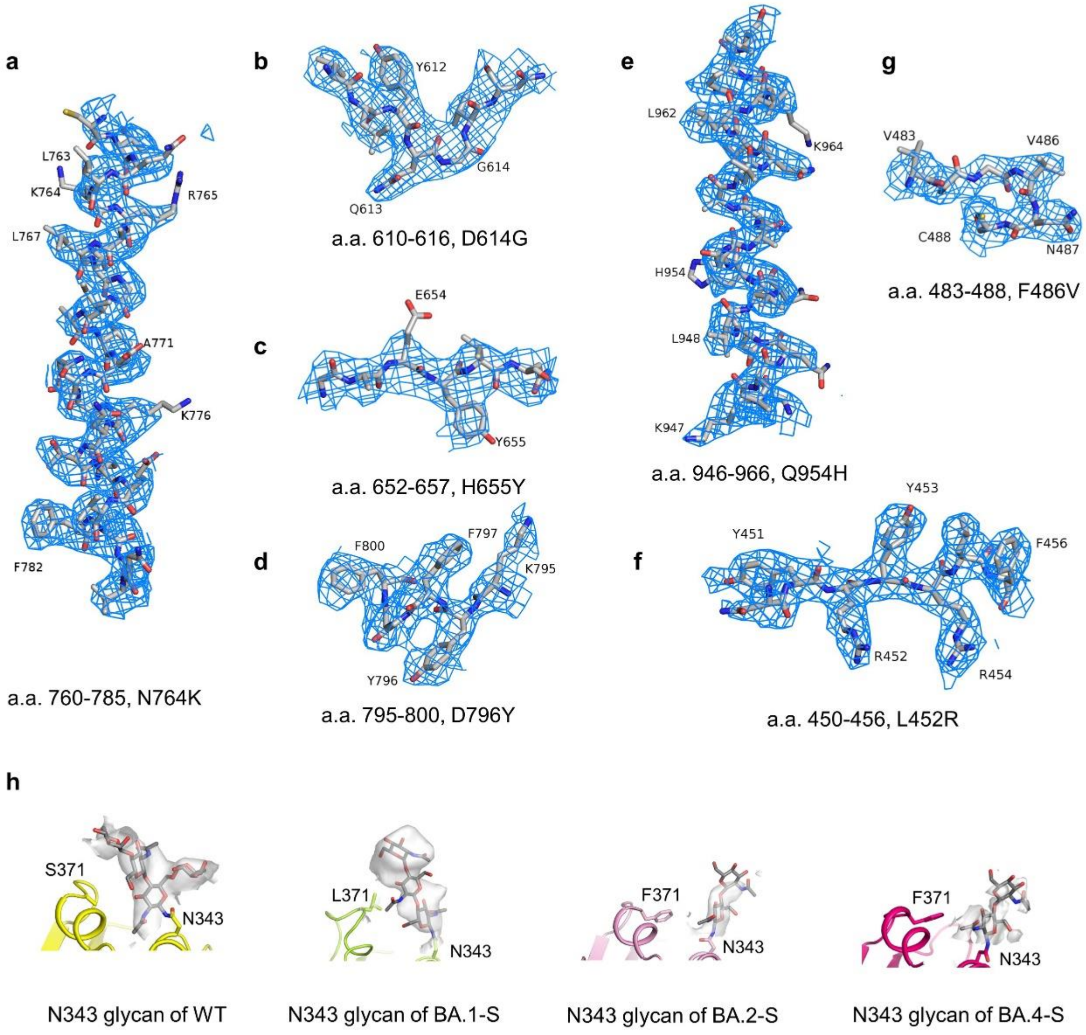


**Supplementary Fig.7**

**Representative cryo-EM density maps.**

**a-c,** Cryo-EM density map of BA.2-SA is shown at threshold of 7 σ. **d-e,** Cryo-EM density map of BA.3-SA is shown at threshold of 7 σ. **f-g,** Cryo-EM density map of BA.4-SA is shown at threshold of 7 σ. **h,** Cryo-EM density map of N343 glycan from WT spike in complex with S309 (6WPT), BA.1-S, BA.2-S, and BA.4-S is shown at threshold of 8 σ, 10 σ, 5 σ, 4 σ, respectively.

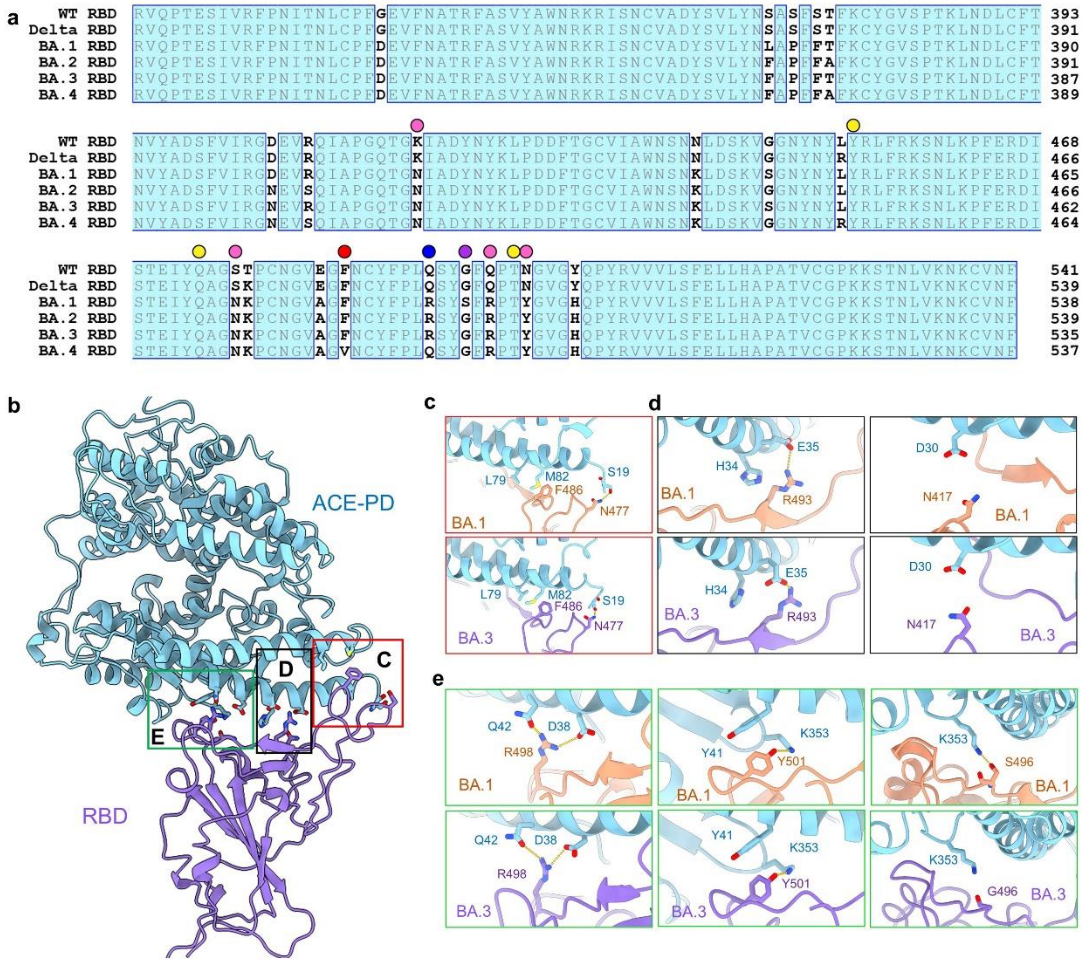


**Supplementary Fig.8**

**Mutations on the interface of RBD from Omicron sub-lineages and ACE2 underlie their enhanced affinity.**

**a,** Sequence alignment of the RBD from indicated SARS-CoV-2 variants. The sequences are aligned using ClustalX. Identical residues are shaded cyan. Residues mapped to the interface with ACE2 are indicated with spheres, yellow for invariant ones and pink for mutated ones. Q493R, indicated by blue sphere, is found in BA.1/2/3. G496S, indicated by purple sphere, is found in BA.1. F486V, indicated by red sphere, is found in BA.4. **b,** Interaction interfaces between RBD and the PD of ACE2. The boxed regions (from right to left) are shown in **c, d** and **e**. **c**, S477N of RBD in BA.1 and BA.3 forms the H-bond with Ser19 of ACE2. **d**, Q493R of BA.1 and BA.3 forms new polar interactions with Glu35 of ACE2 via the salt bridges and K417N disrupts the original interaction with Asp30 of ACE2. **e**, Q498R of BA.1 and BA.3 forms new polar interactions with Asp38 of ACE2 via the salt bridges and

N501Y of BA.1 and BA.3 loses the interaction with Tyr41 of ACE2. G496S of RBD in BA.1 sub-lineages form new H-bond with Lys353 of ACE2.

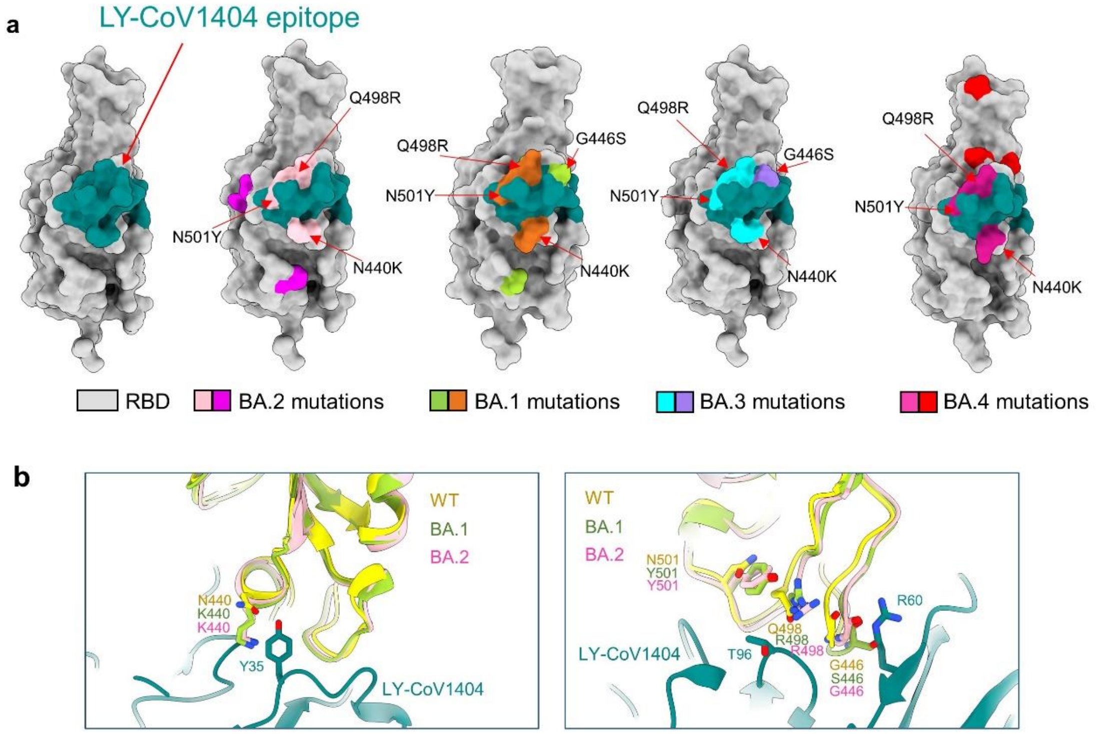


**Supplementary Fig. 9**

**Mapping RBD mutations to representative Class 3 neutralizing antibodies epitopes underlie their neutralizing ability.**

**a,** The binding epitope of antibody LY-CoV1404 bound with the RBD is colored black green (left panel), which overlaps with the mutated residues on BA.2 (middle panel, colored pink), BA.1 (middle panel, colored brown), BA.3 (middle panel, colored cyan), and BA.4 (right panel, colored violet) relative to the RBD (WT) are N440K, G446S, Q498R, and N501Y**.** The mutations shared only by BA.2 and BA.4 are colored magenta (left panel). The mutated residues on BA.1 (middle panel), BA.3 (middle panel) and BA.4 (right panel) different from BA.2 are colored green, purple and red, respectively. **(b)** LY-CoV1404 mapped onto the RBD of BA.2-S.

**Supplementary Table 1**

**Cryo-EM data collection and refinement statistics.**

| | | | | |
|---|---|---|---|---|
| **Data collection** | | | | |
| EM equipment | | Titan Krios (Thermo Fisher Scientific) | | |
| Voltage (kV) | | 300 | | |
| Detector | | Gatan K3 Summit | | |
| Energy filter | | Gatan GIF Quantum, 20 eV slit | | |
| Pixel size (Å) | | 1.087 | | |
| Electron dose (e-/Å2) | | 50 | | |
| Defocus range (μm) | | -1.4 ~ -1.8 | | |
| Sample | BA.2-SA | BA.3-SA | | BA.4-SA |
| Number of collected micrographs | 5,687 | 3,783 | | 5,215 |
| **3D Reconstruction** | | | | |
| Software | | cryoSPARC | | |
| Sample | Overall | 3 "UP" Overall | 2 "UP" Overall | Overall |
| Number of used particles (Overall) | 131,412 | 248,660 | 55,521 | 458,101 |
| Resolution (Å) | 3.3 | 3.4 | 3.8 | 2.8 |
| Symmetry | | C1 | | |
| Map sharpening B-factor (Å2) | 109.1 | 151.2 | 108.2 | 111.3 |
| **Refinement** | | | | |
| Software | | Phenix | | |
| Cell dimensions | | | | |
| a=b=c (Å) | | 313.056 | | |
| α=β=γ (°) | | 90 | | |
| Model composition | | | | |
| Protein residues | 4,786 | 4,786 | 4,153 | 4,786 |
| Side chains assigned | 4,786 | 4,786 | 4,153 | 4,786 |
| Sugar | 104 | 104 | 93 | 104 |
| R.m.s deviations | | | | |
| Bonds length (Å) | 0.003 | 0.003 | 0.005 | 0.003 |
| Bonds Angle (°) | 0.586 | 0.598 | 0.661 | 0.665 |
| Ramachandran plot statistics (%) | | | | |
| Preferred | 93.62 | 93.45 | 92.38 | 93.49 |
| Allowed | 6.21 | 6.21 | 7.32 | 6.17 |
| Outlier | 0.26 | 0.34 | 0.3 | 0.34 |
